# PATRIOT: A pipeline for tracing identical-by-descent chromosome segments to improve genomic prediction in self-pollinating crop species

**DOI:** 10.1101/2020.10.17.343780

**Authors:** Johnathon M. Shook, Daniela Lourenco, Asheesh K. Singh

## Abstract

The lowering genotyping cost is ushering in a wider interest and adoption of genomic prediction and selection in plant breeding programs worldwide. However, improper conflation of historical and recent linkage disequilibrium between markers and genes restricts high accuracy of genomic prediction (GP). Multiple ancestors may share a common haplotype surrounding a gene, without sharing the same allele of that gene. This prevents parsing out genetic effects associated with the underlying allele of that gene among the set of ancestral haplotypes. We present ‘Parental Allele Tracing, Recombination Identification, and Optimal predicTion’ (i.e., PATRIOT) approach that utilizes marker data to allow for a rapid identification of lines carrying specific alleles, increases the accuracy of genomic relatedness and diversity estimates, and improves genomic prediction. Leveraging identity by descent, PATRIOT showed an improvement in GP accuracy by 16.6% compared to the traditional rrBLUP method. This approach will help to increase the rate of genetic gain and allow available information to be more effectively utilized within breeding programs.

## INTRODUCTION

Crop domestication has caused extreme genetic bottleneck, with a reduction in genetic diversity in domesticated crops compared to wild ancestors including in soybean (*Glycine max* L. Merr.) (Hyten et al. 2006). Consequently, the number of ancestral individuals that are represented in modern cultivars is quite low (Gizlice et al. 1994). For example, 17 founding lines contributed 75% of the genes in modern U.S. soybean cultivars, and 95% of genes could be traced to 35 ancestral lines, demonstrating an extremely narrow genetic variation challenging breeding progress. This is not confined to soybean alone, as other crops have similar challenges (Bennett et al. 2012, Smith 2007).

The narrow genetic variability within modern breeding programs is a concern for breeders, as low diversity implies an incomplete sampling of favorable alleles as breeders attempt to improve crop performance and plasticity (Kisha et al. 1998). Furthermore, the likelihood of untapped resistance to biotic and abiotic stresses and unavailability of favorable genes is high (Burdon 2001). Low genetic diversity also negatively influences the response to selection, i.e., genetic gain (Tanksley and McCouch 1997). In soybean, reliance on a single source for soybean cyst nematode (SCN) resistance has increased the ability of SCN to reproduce at a higher rate, necessitating the need to bring in additional sources of resistance (Tylka 2007). Tracking identity-by-descent (IBD) presents unique advantages that can benefit ongoing plant breeding efforts in utilizing the narrow genetic germplasm pool within modern varieties effectively.

Within breeding populations, genes or marker alleles can be expressed as either IBD (individuals share nucleotide sequence; marker allele is the same by inheritance from a shared ancestor) or identical-by-state (IBS) (individuals share nucleotide sequence; marker allele is the same independent of the origin) (Lynch and Walsh 1998). IBD data provides greater information than IBS, as the nucleotide sequence between two adjacent IBD marker alleles from one parent in an individual is also inherited from that same parent at a high probability, barring mutation or double recombination. This physical linkage can extend across multiple loci on a segment of chromosome, depending on local recombination rates; and can be captured using genetic markers. Further, linkage disequilibrium (LD) between a set of markers or gene loci due to physical linkage, mutation, population stratification, or recombination rates may be present (Slatkin 2008). When recombination is low within a region of multiple marker loci, it becomes possible to identify haplotypes, or runs of multiple markers which are consistently inherited together (Daly et al. 2001).

Current genomic selection models effectively model IBS relationships between lines and utilize historic linkage between markers and the trait of interest. This approach has worked reasonably well (Sorrells 2015), which can be attributed to the similarity between IBS and IBD relatedness metrics. In cases where LD is high locally, IBS relationships are more similar to those calculated based on IBD. The use of IBD can improve relationship estimation (Li et al. 2014), kinship and population structure (Morrison 2013), and also is useful for genetic mapping (Dawn Teare and Barrett 2005). The haplotype information from IBD due to inheritance from a recent common ancestor can therefore enable more accurate relationship estimates and improve the effectiveness of genomic selection with IBD-based genomic selection approaches. However, in order to take full advantage of the benefits of IBD data, it is first necessary to track true IBD segments within the population.

The advancements in genetic marker technology have revolutionized the understanding of existing breeding germplasm, allowing the identification of QTL responsible for variation in a trait of interest using genome-wide association or QTL mapping studies, as well as selection of genotypes which contain a QTL of interest in marker-assisted selection (MAS) or marker assisted backcrossing (MABC). Nearly the entire USDA collection of soybean varieties has been genotyped using the SoySNP50K single nucleotide polymorphism (SNP) array containing genome wide markers (Song et al. 2015). Large-scale genotyping efforts have been shown to be useful for scanning germplasm collections in multiple crops (Moellers et al. 2017, de Azevedo Peixoto *et al*. 2017, Yu et al. 2016), and for conducting genome wide association studies using the germplasm collection accessions (Coser et al. 2017, Moellers et al. 2017, Zhang et al. 2017). Alternatively, genome-wide markers can be used to predict performance of untested lines (i.e., genomic prediction (GP)) and subsequently select new varieties with the greatest expected genotypic values (genomic selection (GS)). GS is becoming mainstream in mid-to large breeding programs (Hickey et al. 2017), as it unlocks new opportunities to perform selections in early generations and predict parental suitability (Battenfield et al. 2016, Yao et al. 2018). This leads to the ability to select improved lines reliably with less field testing and speed their re-use as parents in a breeding program. However, GS models require continual model training and validation as a reduction in prediction accuracy has been reported after two to three generations (Jannick 2010). Widespread adoption of GS may require higher prediction accuracy to make up for the additional cost of generating marker data and phenotyping a training population. Therefore, numerous efforts have been made to improve the prediction accuracy of GS models (Jia et al. 2012, Solberg et al. 2008, Habier et al. 2011). One such approach is to better utilize LD information to track chromosome regions inherited from each parent in regions of high LD (Thompson 2013). In this approach, fewer markers are needed for filial generations as so-called “tag” markers can be used to elucidate haplotypes in lines and progenies (Johnson et al. 2001).

While previous efforts have relied on using haplotypes based on observed LD between markers, we explore an alternative approach of tracking the parental source of each allele. Two main distinctions between the approaches should be noted: 1) our approach does not assume any previous evidence of haplotypes or LD, instead utilizing markers which could only have been inherited from exactly one of the direct parents to define IBD segments, and 2) individuals which would otherwise have the same estimated effect from a shared haplotype can now be assigned different estimated effects due to tracking exactly which ancestral line a haplotype was inherited from.

We test an approach hereafter named “Parental Allele Tracing, Recombination Identification, and Optimal predicTion” (PATRIOT) that utilizes raw marker data for tracking IBD inheritance of chromosome segments, enabling the rapid identification of lines carrying specific alleles, increasing the accuracy of genomic relatedness and diversity estimates, and improving genomic prediction and selection performance. Using the SoyNAM population (Song et al. 2017), which includes 39 parents crossed to a common parent and 5176 recombinant inbred lines, we explored the effectiveness of GS with additional information conferred with IBD. We traced chromosome segments from parent to progeny, followed by the calculation of an allelic score for each parental source of each SNP. These allelic scores were used in place of the raw marker data in order to allow the incorporation of IBD data into a GS pipeline.

## MATERIALS AND METHODS

### Pedigree records

Pedigrees for public breeding lines tested in the Uniform Soybean Tests were recorded based on reporting in their last year of testing in the Northern tests (https://www.ars.usda.gov/midwest-area/west-lafayette-in/crop-production-and-pest-control-research/docs/uniform-soybean-tests-northern-region/) or Southern tests (https://www.ars.usda.gov/southeast-area/stoneville-ms/crop-genetics-research/docs/uniform-soybean-tests/). Additional breeding records were recorded from cultivar release papers, primarily from Crop Science (https://acsess.onlinelibrary.wiley.com/journal/14350653), the Journal of Plant Registrations (https://acsess.onlinelibrary.wiley.com/journal/19403496), and Canadian Journal of Plant Science (https://www.nrcresearchpress.com/journal/cjps). Pedigree information for other lines in the NPGS soybean germplasm collection were downloaded from https://npgsweb.ars-grin.gov/gringlobal/search.aspx?. The pedigree information used in this study is provided **Supplemental File 1** and is also available from GitHub (https://github.com/SoylabSingh/PATRIOT).

### Marker data

#### Soybean Nested Association Mapping (SoyNAM) panel

SNP marker data for 5149 SoyNAM RILs, as well as their parents, were downloaded from SoyBase (https://soybase.org/SoyNAM/index.php), using the Wm82.a2 reference genome for downloading. For the SoyNAM panel, 4289 SNP markers were used in the analysis. Markers were re-ordered prior to tracing and imputation based on the composite linkage map created in previous work (Song et al. 2017). The ancestral source of each chromosome segment was identified using the pipeline illustrated in Figure 1.

**Figure 1.**
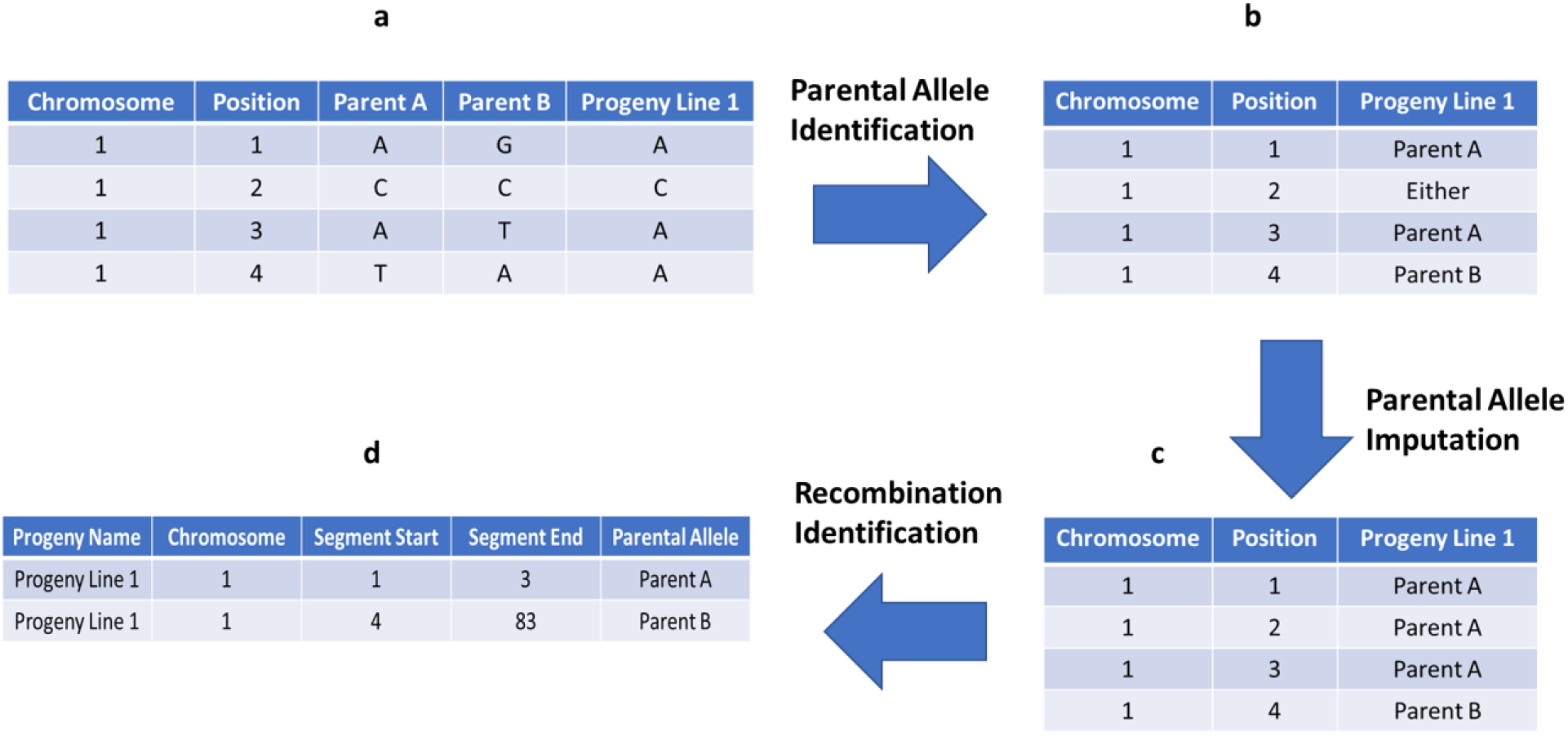
General workflow of Parental Allele Tracing, Recombination Identification, and Optimal predicTion (PATRIOT) input feature preparation for implementation in genomic selection: (a) Raw marker data is provided for both parent and progeny genotypes, b) Parental alleles encoded for those markers which can be conclusively traced to a specific parent, c) Alleles previously not assigned to a specific parent are imputed based on flanking markers, d) Those chromosome segments identical-by-descent from each parent are compiled. The “Position” column refers to the marker order and is provided only for demonstration purposes.

#### Released cultivars and isolines

We used 868 lines that were the progeny of parental lines, and wherein both parent and progeny were genotyped with the SoySNP50k SNP set. Near-isogenic lines derived from backcrossing schema were also included in this pipeline, which may make some statistical metrics not applicable for this panel. SNP marker data for all accessions in the GRIN database were downloaded from Soybase.org (https://soybase.org/dlpages/#snp50k) as a VCF file, with positions annotated based on the Wm82.a2 reference genome. Pre-processing to remove SNPs aligned to scaffolds or the mitochondria left 42,080 SNP markers aligned to the Wm82.a2 reference genome and used in further analysis. Missing SNP data were imputed using Beagle 4.0 with default settings (Browning and Browning 2007). This panel will be referred to as the “868/50K panel” for brevity.

### Performance data

Phenotypic records for the SoyNAM recombinant inbred line mapping population were downloaded from SoyBase (https://soybase.org/SoyNAM/index.php). Replicated entries’ phenotypic records from within a single environment were used to calculate BLUPs for those lines, while unreplicated entries were incorporated using the raw phenotypic values. The “Corrected Strain” column was used to connect phenotypes with genotypic records. Phenotypic records were available from 2011 (IL and NE), 2012 (IA, IL, IN, KS, MI, MO, NE, OH^1^, and OH^2^), and 2013 (IA, IL, IN, KS, and MO).

*Phytophthora* root rot resistance ratings were queried from the National Plant Germplasm System (https://npgsweb.ars-grin.gov/gringlobal/search.aspx) for each of the ancestors of the modern cultivar “Rend” (Nickell et al. 1999). “Rend” was selected for demonstration of the multi-generation chromosome segment tracing code due to both parents and all four grandparents being genotyped with the same platform, as well as the major resistance gene segregating within the pedigree.

### PATRIOT workflow and code development

PATRIOT workflow utilizes LD and haplotype in a novel way to improve genomic prediction. Specifically, this system allows for the tracing of chromosomal segments from the immediate parents to the offspring, and also to trace chromosomal segments through multiple generations. The allele tracing code outputs can be used as inputs into a modified genomic selection code, wherein the nominal class data are converted to numeric through the use of differences from the mean. Custom R scripts were developed to identify SNPs which could only come from one of the listed parents (hereafter “anchor markers”, **Figure 1a, 1b**), followed by imputation of SNPs of fixed markers based on surrounding anchor markers (**Figure 1c**). Code for identifying anchor markers, imputation, multi-generation tracing, and recombination zone identification are available as **Code 1**, **Code 2**, **Code 3**, **Code 4**, respectively (**Supplementary File 2**). Genomic prediction was evaluated using rrBLUP in R with raw marker data and allele tracing alternatives (**Code 5**) (**Supplementary file 2**).

### Chromosomal tracing and Identity By Descent (IBD)

As a proof of concept, tracing of chromosome segment inheritance within the pedigree of soybean cultivar “Rend” was performed. After ensuring consistency between expected results and the outputs, chromosome tracing was performed on the remainder of the 868/50K panel. Following completion of the single-generation tracing pipeline, the multi-generation tracing script was run on traced lines to allow visualization of multiple generations of inheritance and recombination.

In addition to the 868/50K panel, SoyNAM project parents and RILs were investigated with the chromosome tracing pipeline. The A/B genotype representation data available from SoyBase was utilized to impute chromosomal segments. Even with a sparse marker coverage, recombination events were still identifiable (**Supplemental File 2**).

### Genomic prediction models

To expand on the usefulness of the chromosome tracing pipeline outlined in **Figure 1**, we used the SoyNAM panel to evaluate accuracy of genomic prediction using ancestral alleles. Genomic prediction was evaluated for multiple traits using the 39 SoyNAM RIL populations based on the phenotypic records available from the SoyNAM project and all 4289 available markers. All comparisons were made using 80% of individuals for training and predicting on the remaining 20% of individuals. For each marker, an allelic differential estimator (ADE) was calculated as:

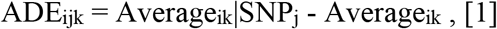

for environment i, marker j, and trait k. The ADE value therefore is not regressed towards the mean to account for multiple regression. Instead, these values replace the marker representation as an input to GS models that evaluate the performance of this new approach (**Table 1**). They allow for the use of many distinct ancestral haplotypes in linear regression-based models based on the sign and relative scale of the estimated haplotype effect.

Traditional rrBLUP performance was evaluated using mixed.solve, a function in “rrBLUP” (Endelmann 2011). The rrBLUP-PATRIOT analysis was performed using mixed.solve, but replacing the marker input data (0,1,2) with a matrix of ADEs calculated in PATRIOT. The mean observed phenotype of lines with top 10% of predicted performance using rrBLUP and PATRIOT were compared, as well as the difference in phenotype between selected lines and the base population. For yield, five-fold cross-validation was used to reduce sampling bias in the estimation of GP accuracy for each method.

**Table 1.** Simplified matrix showcasing 5 potential progeny, parents, and their ADEs. ADEs were calculated using the full panel (more than one family) and with unequal population sizes, so marker effects are not necessarily equal and opposite. Progeny 2 and 3 differ based on the site of recombination around SNP 3. Progeny 4 represents the optimal combination of ancestral alleles available from within that population, but not necessarily within the full panel. Progeny 5 represents the global optimum progeny from within the panel; however, this progeny would require multiple crosses to introgress segments from all three parents. ADE scale is shown based on potential values for yield in terms of kg ha^−1^ in soybean.

|  | SNP 1 | SNP 2 | SNP 3 | SNP 4 | SNP 5 | SNP 6 | SNP 7 |
| --- | --- | --- | --- | --- | --- | --- | --- |
| Parent 1 | 45 | -23 | 70 | 14 | -56 | 73 | 15 |
| Parent 2 | -40 | 17 | -65 | -50 | -15 | -51 | 70 |
| Parent 3 | -53 | 20 | -71 | 106 | 69 | -36 | -43 |
| Progeny 1 | 45 | -23 | 70 | -50 | -15 | -51 | 70 |
| Progeny 2 | -40 | 17 | -65 | 14 | -56 | 73 | 15 |
| Progeny 3 | -40 | 17 | 70 | 14 | -56 | 73 | 15 |
| Progeny 4 | 45 | 17 | 70 | 14 | -15 | 73 | 70 |
| Progeny 5 | 45 | 20 | 70 | 106 | 69 | 73 | 70 |

## RESULTS

### Recombination Identification

For the 868/50Kpanel, 13.14% of all SNPs were unassigned to a specific parent. For the SoyNAM panel, 6.78% of all SNPs were unable to be assigned to a specific parent. Using the SoyNAM panel marker data after PATRIOT IBD tracing and imputation, we examined the rates of recombination throughout the genome. Of the 5149 RILs examined, we found total recombinations per line ranged from 10 to 557, with an average of 50.9 recombinations per line. 18808/102960 (18.3%) chromosomes were inherited intact from one parent or another. 5011 RILs inherited at least one intact chromosome from a parent.

### Chromosomal Segment Tracing and Recombination Events

Chromosomal segments were traced in the 868/50K panel using the PATRIOT framework. To demonstrate the PATRIOT workflow, we trace the inheritance of the major *Phytophthora* root rot (PRR) resistance locus *Rps1* (**Figure 2**). Williams 82 (i.e., PI518671) inherited the *Rps1k* allele (that confers PRR resistance), as a long introgression (shown in green) on chromosome 3 from Kingwa (i.e., PI548359). This allele is then transmitted from Williams 82 to Resnik (i.e., PI534645) in a smaller chromosomal segment around *Rps1k*. However, the resistance allele was not passed on to Resnik’s progeny, Rend (i.e., PI606748). Resnik is therefore more suitable than Rend to breed for Phytophthora resistance. Chromosomal tracing over multiple generations allows presence/absence characterization for the *Rps1k* allele without the need for allele-specific markers and can reduce the need for phenotyping in disease nurseries, as allele state is known by virtue of IBD. **Figure 2** gives a visual chromosomal segment tracing that is applicable to all varieties with available pedigree records that have been genotyped.

Recombination events can be visually identified when examining multiple generations within **Figure 2** (or similar plots) in two ways using the chromosome 3 example: (i) between Williams 82 and Resnik, the length of the green segment surrounding *Rps1k* is greatly reduced in Resnik, indicating recombination during the cross of Asgrow 3127^4^ x Williams 82, and (ii) a segment of the soft red “AmbiguousParentage” class appears in the progeny, which indicates that recombination occurs somewhere within this region, but could not be delimited between two adjacent markers due to multiple markers being alike by state in the parents. This occurs in Asgrow 3127 (i.e., PI556511) on chromosome 3, separating large segments inherited from Williams and Essex.

While the *Rps1k* example is provided, the PATRIOT framework is applicable to trace chromosomal regions and for IBD characterization of important genes through generations, as well as to visualize nearby recombination events.

**Figure 2.**
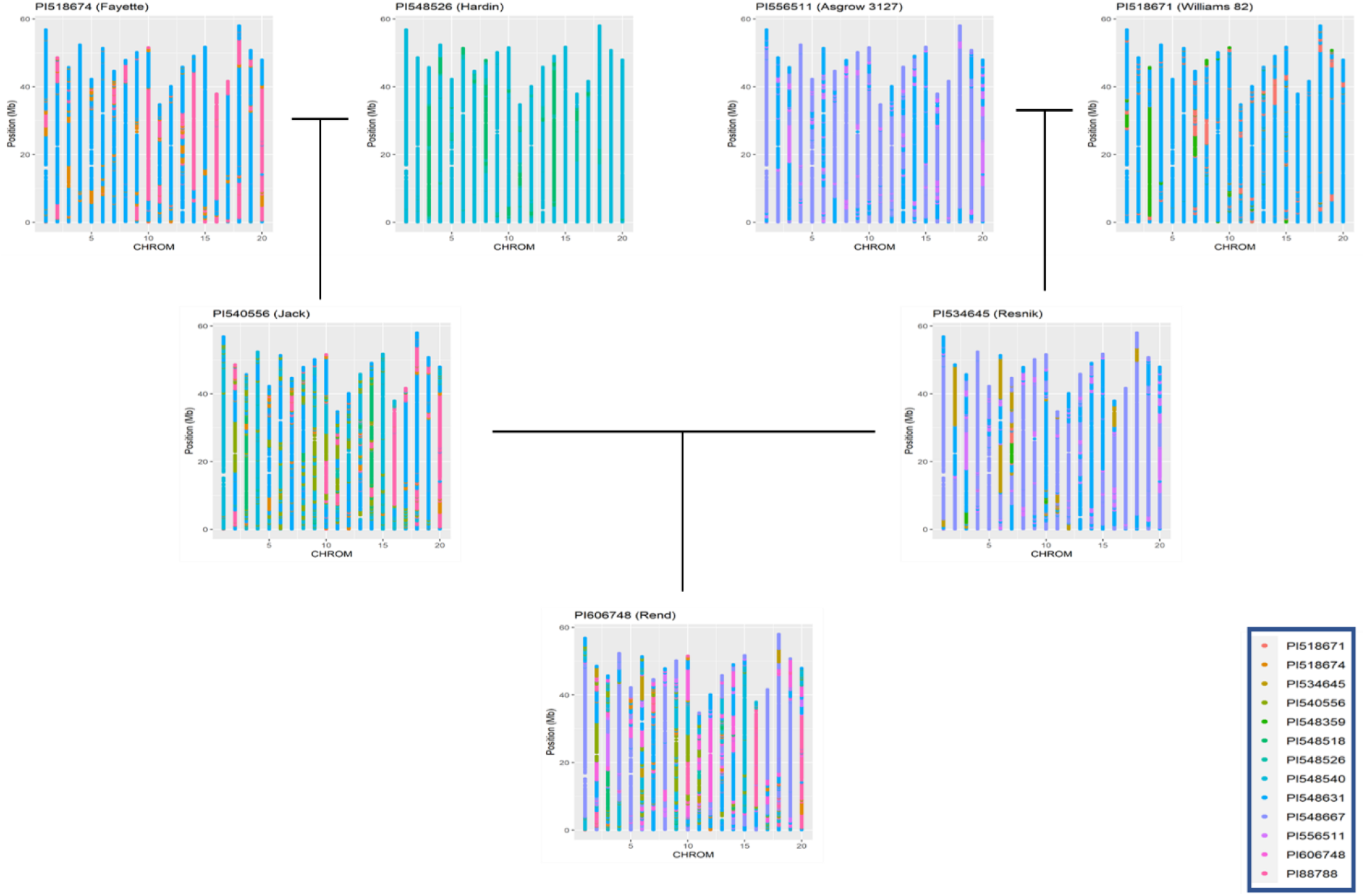
Scatterplot maps of chromosome segments inherited from ancestral sources, traced through progenitors of soybean cultivar Rend (i.e., PI606748). Chromosome number (based on Wms82.a2 reference genome) is plotted left to right on the x-axis, while position is plotted on the y-axis.

### Comparison of genomic prediction accuracy using SoyNAM

To examine the relative effectiveness of rrBLUP with PATRIOT (PATRIOT GS) compared to traditional rrBLUP (rrBLUP GS), yield predictions for 16 environments from each model were generated using the same randomized testing set for each model (Table 2). Results from the two GS approaches are presented in **Table 2**. A 16.6% increase was attained in genomic prediction accuracy by using PATRIOT GS compared with traditional rrBLUP (0.557 vs. 0.478). Using a scenario of selecting 10% (and discarding 90%) from the SoyNAM RIL population and comparing to the overall SoyNAM RIL population mean, PATRIOT GS had a 8.6% greater selection differential among the selected RILs over basic rrBLUP GS (an increase of +538.7 in PATRIOT GS vs +496.1 kg ha^−1^ in rrBLUP GS).

**Table 2.**
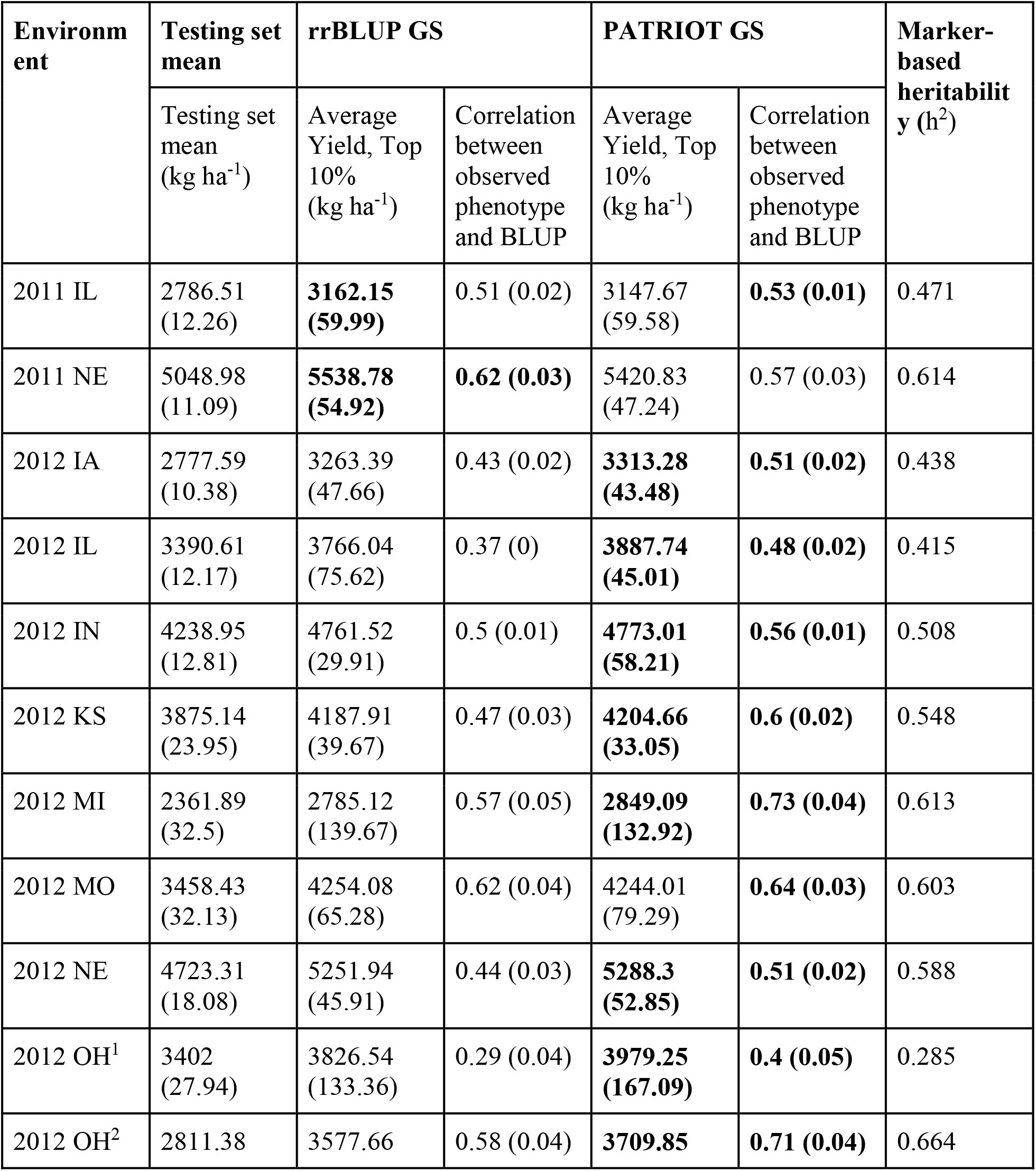
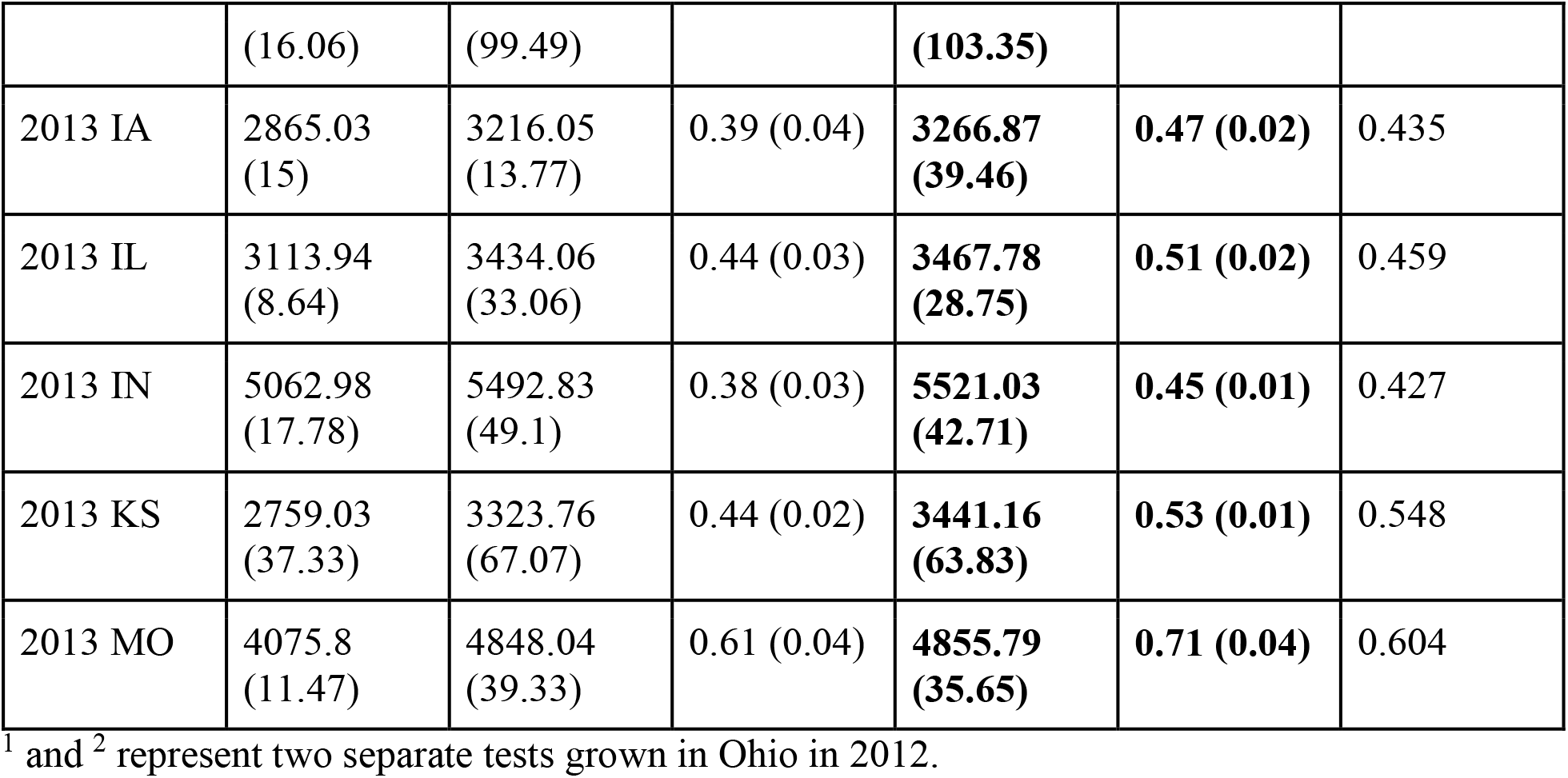
Comparison of the effectiveness of genomic selection methods rrBLUP GS and PATRIOT GS for yield. In each environment, the best model for each metric is highlighted in bold. Standard deviation given in parentheses. Each row contains results from a single site/year combination; “2011 IL” is therefore from an experiment in Illinois during 2011.

| Environm<br>ent | Testing set<br>mean | rrBLUP GS | | PATRIOT GS | | Marker-based<br>heritabilit<br>y ($h^2$) |
| --- | --- | --- | --- | --- | --- | --- |
|  | Testing set<br>mean<br>(kg ha<sup>-1</sup>) | Average<br>Yield, Top<br>10%<br>(kg ha<sup>-1</sup>) | Correlation<br>between<br>observed<br>phenotype<br>and BLUP | Average<br>Yield, Top<br>10%<br>(kg ha<sup>-1</sup>) | Correlation<br>between<br>observed<br>phenotype<br>and BLUP |  |
| 2011 IL | 2786.51<br>(12.26) | **3162.15**<br>**(59.99)** | 0.51 (0.02) | 3147.67<br>(59.58) | **0.53 (0.01)** | 0.471 |
| 2011 NE | 5048.98<br>(11.09) | **5538.78**<br>**(54.92)** | **0.62 (0.03)** | 5420.83<br>(47.24) | 0.57 (0.03) | 0.614 |
| 2012 IA | 2777.59<br>(10.38) | 3263.39<br>(47.66) | 0.43 (0.02) | **3313.28**<br>**(43.48)** | **0.51 (0.02)** | 0.438 |
| 2012 IL | 3390.61<br>(12.17) | 3766.04<br>(75.62) | 0.37 (0) | **3887.74**<br>**(45.01)** | **0.48 (0.02)** | 0.415 |
| 2012 IN | 4238.95<br>(12.81) | 4761.52<br>(29.91) | 0.5 (0.01) | **4773.01**<br>**(58.21)** | **0.56 (0.01)** | 0.508 |
| 2012 KS | 3875.14<br>(23.95) | 4187.91<br>(39.67) | 0.47 (0.03) | **4204.66**<br>**(33.05)** | **0.6 (0.02)** | 0.548 |
| 2012 MI | 2361.89<br>(32.5) | 2785.12<br>(139.67) | 0.57 (0.05) | **2849.09**<br>**(132.92)** | **0.73 (0.04)** | 0.613 |
| 2012 MO | 3458.43<br>(32.13) | 4254.08<br>(65.28) | 0.62 (0.04) | 4244.01<br>(79.29) | **0.64 (0.03)** | 0.603 |
| 2012 NE | 4723.31<br>(18.08) | 5251.94<br>(45.91) | 0.44 (0.03) | **5288.3**<br>**(52.85)** | **0.51 (0.02)** | 0.588 |
| 2012 OH<sup>1</sup> | 3402<br>(27.94) | 3826.54<br>(133.36) | 0.29 (0.04) | **3979.25**<br>**(167.09)** | **0.4 (0.05)** | 0.285 |
| 2012 OH<sup>2</sup> | 2811.38 | 3577.66 | 0.58 (0.04) | **3709.85** | **0.71 (0.04)** | 0.664 |
|  | (16.06) | (99.49) |  | **(103.35)** |  |  |
| 2013 IA | 2865.03<br>(15) | 3216.05<br>(13.77) | 0.39 (0.04) | **3266.87**<br>**(39.46)** | **0.47 (0.02)** | 0.435 |
| 2013 IL | 3113.94<br>(8.64) | 3434.06<br>(33.06) | 0.44 (0.03) | **3467.78**<br>**(28.75)** | **0.51 (0.02)** | 0.459 |
| 2013 IN | 5062.98<br>(17.78) | 5492.83<br>(49.1) | 0.38 (0.03) | **5521.03**<br>**(42.71)** | **0.45 (0.01)** | 0.427 |
| 2013 KS | 2759.03<br>(37.33) | 3323.76<br>(67.07) | 0.44 (0.02) | **3441.16**<br>**(63.83)** | **0.53 (0.01)** | 0.548 |
| 2013 MO | 4075.8<br>(11.47) | 4848.04<br>(39.33) | 0.61 (0.04) | **4855.79**<br>**(35.65)** | **0.71 (0.04)** | 0.604 |

<sup>1</sup> and <sup>2</sup> represent two separate tests grown in Ohio in 2012.

To help explain the cause of the difference in performance improvement between genomic prediction accuracy (+16.6%) and genomic selection effectiveness (+8.6%) (both compared to rrBLUP), we further examined the yield predictions from the 2012 OH^1^ environment, which showed a large increase in GP accuracy (+39.5%) but only slight increase in genomic selection effectiveness (+3.8%). When examining the bottom 10% of predicted lines (rather than top 10% as before), the genomic selection effectiveness was 52.7% greater using PATRIOT than rrBLUP. This finding, coupled with smaller average absolute error terms using PATRIOT, suggests that the GP accuracy increase came from decreased error terms throughout the full range of phenotypes, allowing for better rankings. Indeed, using a 5% selection level for high GEBVs using PATRIOT resulted in a 29.8% increase in average observed phenotype compared to rrBLUP in the 2012 OH^1^ set.

## DISCUSSION

Some of the earlier efforts in soybean chromosomal tracing involved RFLP markers, as researchers traced chromosome segments in 67 genotypes through generations (Lorenzen 1994). The transition to SNP markers as more mainstream marker technology enables better genome coverage to trace chromosomal segments from progenitors (Letcher and King 2001), with increased resolution for recombination identification (Yu et al. 2011). However, the biallelic nature of SNP markers is a limitation for more refined haplotype generation. In the 868/50K panel, 13.14% of all markers could not be definitively traced back to their ancestral source. While some portion of this unassigned group can be attributed to heterozygous allele state in either one of the parents or the progeny, a substantial portion is due to recombination in the affected area in which both parents are IBS at several consecutive markers. Less singletons were identified in the SoyNAM panel, possibly due to the non-identification of small double-recombination events, as well as due to the use of a linkage map for marker ordering, which by its nature reduces the likelihood of mis-ordering markers along a chromosome or linkage group. The genome tracing of large segments through multiple generations enables breeders to follow genes of interest throughout the pedigrees of modern lines (Bruce et al. 2020). This allows for a rapid identification of lines containing the desired allele even if allele specific markers are not available. Visualization of relatedness of lines based on IBD metrics similar to **Figure 2** allows breeders to rapidly identify pairings of lines with high genetic diversity as parents to create breeding families (Liu & Anderson 2003).

While IBD can be traced in many released public cultivars on the basis of markers from the SoySNP50K chip in soybean, applicability to breeding programs during the development of new pure lines requires a cost-effective genotyping system to allow genotyping of these lines at an earlier stages of development. This can be achieved by utilizing a smaller, less expensive genotyping array such as the SoyNAM6K BeadChip (Song et al. 2017) to genotype experimental lines.

The PATRIOT framework facilitates the identification of lines for breeding purposes that have favorable genes linked in coupling, as well as in situations where breaking the linkage drag is imperative. For example, SCN resistance from PI494182 was determined to carry a risk of linkage drag (St-Amour et al. 2020). Likewise, SCN resistance from the commonly used donor PI88788 was initially associated with considerable linkage drag (Cregan et al 1999). With the use of PATRIOT, parents can be readily identified which contain the gene(s) of interest with the least amount of additional introgressed region(s), thereby reducing the likelihood of linkage drag, and concurrently deploy it in a GS pipeline. With an additional generation of traced progeny, those regions negatively associated with another trait can be identified to inform marker based decisions.

Much like genome-wide association studies (GWAS), genomic prediction and genomic selection models rely on the association between markers and the phenotype of interest. However, the association between marker and phenotype decays in subsequent generations, leading to reduced accuracy without re-training of the model (Habier et al. 2007, Hayes et al. 2009, Jannick 2010). With the chromosome tracing approach, the linkage between marker and phenotype withstands this shift, since parental allele representation is directly incorporated into the marker data. The prediction accuracy decays much more slowly with chromosome tracing because the linkage between marker and phenotype decays only as recombination between marker and underlying gene(s) occurs. Furthermore, multi-generation tracing allows the preservation of information on lineage-specific marker association which can better model the differences in genes linked to a particular marker or set of markers.

The widespread use of PATRIOT GS would be encouraged by the establishment of a fully connected pedigree (fully known relationships between all germplasm utilized) and development of base population resources with equal and ample representation of each parental source within the breeding pool. For example, while the SoyNAM panel can be readily used as a training set for materials derived from any combination of the 40 parents, it’s efficacy is limited to that context, with the exception of including a small number of the parents’ ancestors within the pool. Instead, in some situations, breeding applications would benefit from the development of fully inter-related populations derived from the founder lines, such as through MAGIC design (Xiao et al 2013, Matteo et al 2015) or a NAM population created with founder parents (Yu et al. 2008) that can happen in different crossing cycles. Moreover, most breeding programs have an inherent nested design especially when a few superior parents are used extensively in the development of breeding populations, therefore this effort is not incremental.

The multi-generation chromosome segment tracing aspect of PATRIOT can also be used as a tool to connect QTL mapping studies among related populations. In addition to tracing chromosomal regions within a pedigree, this framework can be used to connect linkage mapping studies using related lines as parents by tracing QTL regions identified in related parents in separate studies to their ancestral sources. This allows for a meta-analysis to utilize the increased power which comes from having multiple mapping populations with common ancestry to map marker-trait associations.

However, there are challenges to the PATRIOT framework. In crosses where parents share large runs of IBS or IBD based on marker data, it is difficult to determine which parent is contributing each allele to the progeny. However, if these runs are IBD, the effect on allele estimation is equivalent, regardless of which parent is assigned to the allele. Additionally, a surprising number of singleton marker calls suggests that either double recombination is occurring at a much higher rate than previously believed, or that the reference genome assembly order does not agree with the true marker order. Increased marker density can overcome some of these challenges. Likewise, uncertain regions can be assigned new allele effect classes. For example, Williams 82 (PI518671) has 3,399 out of 42,080 markers which could not be assigned with certainty to a specific parent (Williams or Kingwa). To circumvent this challenge, each of these markers was assigned a new parent class of “PI518671” when tracking segments passed on to progeny, but continue to use ADEs based on the average ADE of parents Williams and Kingwa when predicting its own performance.

PATRIOT genomic prediction accuracy for yield using all populations was greater than the calculated marker-based heritability of the trait in 13 of 16 environments (**Table 2**), suggesting that genomic prediction using ancestral allele tracing can perform better than traditional genomic prediction. Generating separate prediction models in this way for each environment can also be used to reduce the number of environments needed for phenotypic evaluation, as the prediction accuracy very nearly reaches the heritability of the trait itself. The fact that this high level of prediction accuracy was possible with a 6K SNP chip in the SoyNAM populations suggest significant potential cost savings, as the cost of genotyping at this density is less expensive than growing and phenotyping in replicated field plots (Xu et al. 2020). More generally speaking, if small arrays are to continue to be used in community research projects, the array needs to be carefully designed to provide adequate coverage throughout the genome. Consideration of both linkage distance and optimal SNP selection in genic regions should be made a priority. Alternatively, other genotyping platforms such as genotyping-by-sequencing (GBS) can be used to implement this approach, which is able to decrease the negative impact of missing data that is common from GBS (Gardner et al. 2014).

While our genomic prediction models utilized only the immediate parents for calculating allele effect estimates, it is possible to expand the method by combining with the multigeneration IBD tracing script. This approach has an added benefit of bridging the gap between populations that do not share a direct parent but share ancestors in previous generations. By doing so, an increased number of lines can be used for allele effect estimation, further improving the accuracy of these values.

IBD-based genomic selection has the clear potential to improve selection accuracy over existing genomic selection approaches. However, there is a tradeoff due to the significant increase in computational time. While the chromosome segment tracing portion of the workflow need only be run once for any particular genotype, the ADE matrix must be calculated separately for each trait and environment. Fortunately, this calculation can be parallelized, and only needs to be performed for the training population. Typical computation time on an AMD Ryzen Threadripper 1950X for ADE matrix calculation was on the order of one minute without parallelization of the code, while the genomic prediction itself took on the order of three minutes for a dataset with 2500 individuals and 4289 markers. Computation time for the tracing and imputation of alleles within the SoyNAM study totaled 7 hours 41 minutes. However, minor modifications to run each chromosome in parallel on a different computational thread can reduce the wall time to around 35 minutes.

## CONCLUSION

PATRIOT provides a framework for identifying, tracking, and applying IBD information in order to increase effectiveness of genomic selection. Tracking IBD with PATRIOT enables pedigree-based gene tracking through generations, which can be useful for parental selection, as well as for predicting phenotypes for monogenic and oligogenic traits. Relatedness metrics within breeding populations can also be improved due to the specification of IBD allele sharing rather than IBS. The IBD information also works to improve genomic prediction and selection results. This improvement was shown in first-cycle genomic prediction, but should provide additional benefits in later cycles due to the donor-specific allele effect estimation, which does not suffer from the problem of population shift between training and testing sets. The large and consistent benefit shown suggests that chromosome tracing is a quick and efficient way to increase the accuracy of genomic selection models, with no additional cost beyond modestly increased computational time.

## Contributions

JMS conceptualized the project with AKS; JMS conducted the statistical analysis with suggestions from AKS and DL. JMS and AKS prepared the first draft. All authors contributed in writing and editing the manuscript.

## Acknowledgements

Authors sincerely appreciate inputs from Dr. David Grant (USDA-ARS, retired), and Dr. Rex Nelson (USDA-ARS) for assistance with pedigree compilation and suggestions on potential applications for the method. We thank Dr. Kulbir Sandhu, Sarah Jones, and Liz van der Laan for reviewing the manuscript draft.

## Pedigree and Code availability

https://github.com/SoylabSingh/PATRIOT

## List of abbreviations used

GP: Genomic prediction
GS: Genomic selection
rrBLUP: Ridge regression best linear unbiased predictors
PATRIOT: “<u>P</u>arental <u>A</u>llele <u>T</u>racing, <u>R</u>ecombination <u>I</u>dentification, and <u>O</u>ptimal predic<u>T</u>ion”
IBD: Identical-by-descent
IBS: Identical-by-state
LD: Linkage disequilibrium
QTL: Quantitative trait locus
MAS: Marker-assisted selection
USDA: United States Department of Agriculture
NAM: Nested Association Mapping (population)
SNP: Single nucleotide polymorphism

